# Toward space-optimal approximate colored indexes at scale

**DOI:** 10.64898/2026.11.05.764633

**Authors:** Tuna Okçu, Can Alkan, Antoine Limasset

## Abstract

Searching current petabase-scale sequence repositories requires indexes that combine compact storage with high query throughput. Although exact colored indexes retain complete sequence information, their construction and storage costs can become substantial for highly diverse collections. Indexes based on approximate membership query (AMQ) filters offer a compact alternative with fast construction and queries. Being hash-based, these indexes support different seeding schemes without structural changes, while sampling can reduce storage requirements, construction time and query time by an order of magnitude.

Within the AMQ framework, querying many independent filters causes scattered memory accesses, reducing throughput. This motivates interleaved filters, which address this weakness by storing filter slots contiguously by position, enabling cache-friendly access and simultaneous membership queries. However, existing implementations face a trade-off between guaranteed construction and space-efficiency.

Here, we introduce a new indexing framework based on interleaved ZOR filters, fingerprint-based structures with guaranteed construction that require only approximately 1% more space than the theoretical minimum for AMQ. To exploit redundancy between genomes, we introduce a color-preserving merging mechanism that extends fingerprints with membership bits, allowing multiple genomes to share a filter while retaining their individual identities. Together with a lightweight, sketch-based layout planner, this mechanism yields a compact index structure that accommodates collections with widely varying genome sizes and degrees of similarity. Our third contribution extends the AMQ framework beyond contiguous *k*-mers through a low-density minimizer scheme alongside efficient implementations of sensitive seeds, including gapped *k*-mers and randstrobes.

We implement these methods in Zindex, a parallel, SIMD-accelerated Rust index that scales to millions of genomes while supporting diverse seeding and sampling schemes. Zindex indexes the latest RefSeq release of 564,401 assembled genomes in under 22 hours, producing a 289.5 GB index. It further scales to all 2,763,297 genomes of AllTheBacteria, producing a 406.3 GB index in four days with a peak memory footprint of 43.8 GB.

**Availability:** Zindex is available at https://github.com/Malfoy/Zindex

## 1 Introduction

Despite advances in artificial intelligence, including models trained directly on genomic sequences [1,2], the semantics of DNA remain largely unknown. Comparing sequences therefore remains the foundation of biological interpretation, as similarities and differences reveal relationships, conserved functions, and variation [3,4]. In this sense, sequence comparison is the atomic task of sequence bioinformatics, and alignment provides a well-established formulation [5] for which modern algorithms offer highly optimized solutions [6,7].

As individual sequencing experiments reach terabases [8], the Sequence Read Archive (SRA) approaches 100 petabases [9], and both sequencing output and the number of available genomes continue to grow [10], repository-scale resources that make these collections searchable, such as MetaGraph [11] and Logan [12], become increasingly essential. However, even the most efficient alignment algorithms are already costly for individual genomes spanning gigabases [13] and become prohibitively so at repository scale.

This limitation motivates alignment-free methods [14], which scale to massive sequence collections by bypassing explicit alignment to estimate sequence similarity efficiently using rapid set and counting operations on sets of shared short *k*-length substrings (*k*-mers) [15,16,17]. Owing to their scalability and conceptual simplicity, *k*-mer methods have become ubiquitous across bioinformatics, driving core tasks such as assembly [18] and read mapping [19], taxonomic classification [20] and abundance estimation [21], transcript quantification [22,23], variant analysis [24] and copy-number analysis [25], and pangenome analysis [26].

To integrate multi-dataset analyses with minimal redundancy, exact colored de Bruijn graphs (Cortex [27]) store shared *k*-mers once, along with their dataset origins. This family of methods further reduces space by leveraging the sequential nature of genomes—where overlapping *k*-mers form long continuous sequences rather than independent keys—using compacted graphs and sequence-aware indexes [28,11,29,30,31]. However, these representations derive much of their compactness from consecutive *k*-mers, making it less straightforward to exploit sparse sampling schemes such as minimizers [32]. Moreover, building a global graph over the whole collection becomes costly in time and memory at scale, especially with diverse collections and with data that fragment the graphs, such as transcriptomes, metagenomes and variable pangenomes [33,34,35]. Approximate colored indexes, on the other hand, are more flexible, with the potential to efficiently index and query both diverse and redundant data. These approaches represent each *k*-mer set approximately as a Bloom filter [36], a hashing-based structure with membership queries that occasionally return false positives. This simple structure supports fast construction with low memory, making it straightforward to scale indexing by processing datasets independently in parallel. Within this framework, the Sequence Bloom Tree (SBT) [37] uses a hierarchy of Bloom filters, supporting thresholded query semantics: a dataset is considered a match to a query sequence only if the fraction of the query *k*-mers it contains meets a threshold. Even if individual *k*-mer membership queries have relatively high false-positive rates, requiring enough matches within the same dataset can preserve strong sequence-level specificity even for short sequences. Subsequent SBT variants improve on this basic design, most exploiting redundancies between filters for increased compression and faster queries [38,39,40].

Complementing these approaches, the interleaved Bloom filter (IBF) [41,42] stores each bit position contiguously across all datasets in a slice, rather than storing separate filters per dataset. This layout allows each hash function to map a query key directly to a single slice, where bitwise operations evaluate membership across the entire collection simultaneously, improving cache locality and leveraging word-level parallelism. Beyond presence and absence, Needle [43] and REINDEER2 [44] encode approximate abundances using filter-based indexes, enabling quantitative sequence queries across datasets. Dynamic variants also support incremental updates, allowing new datasets to be added without rebuilding the entire index [45]. However, a pure interleaved index requires uniform filter sizes and hash functions across datasets with disparate cardinalities, inflating the space required to accommodate the largest dataset. To reduce this waste, later tools aim to balance intra-IBF cardinalities: COBS [46] groups datasets of similar size into separately sized blocks, while the hierarchical interleaved Bloom filter (HIBF) [47] splits large datasets and merges small ones into a hierarchy of IBFs, which also improves query throughput. Separately, kmindex [48] and PAC [49] partition interleaved matrices by minimizer for cache locality. To determine how datasets should be split and grouped across its hierarchy, the HIBF uses dynamic programming [47].

However, this layout optimization can become costly at large scales, and the resulting hierarchy stores *k*-mers at multiple levels. Beyond layout optimization, hash-based filters can reduce resource requirements by representing each *k*-mer with its minimizer [32], indexing and querying these representatives rather than the full *k*-mers. A *k*-mer is then considered potentially present when its minimizer is found, which preserves true matches but can introduce additional false positives when an alien *k*-mer shares its minimizer with sequences present in the dataset. The SeqAn-based tools Raptor [50] and Needle [43] implement this strategy to reduce storage requirements, construction time, and query time. More recent low-density schemes, such as the mod-minimizer [51] and open-closed mod-minimizer [52], can select fewer representatives for a given window size, offering further gains in scalability.

The flexibility of hash-based filters also extends to seeding, with the SeqAn-based tools supporting gapped *k*-mers [53], which improve sensitivity to substitutions but rely on fixed positional patterns that are vulnerable to insertions and deletions.

With the growing use of long-read sequencing, this limitation motivates moving to seeds such as strobemers [54], which improve robustness to indels. Overall, few indexes exploit the full flexibility of indexing hashes, which allows arbitrary sequence-derived objects to be represented without changing the underlying index structure.

The choice of underlying membership filter offers another opportunity to reduce space, as most approximate collection indexes still use Bloom filters despite the availability of more compact static XOR-based alternatives [55,56]. Taxor [57] demonstrates how these alternatives can be used in an interleaved index, but their probabilistic construction can require retries that complicate scaling to large collections. ZOR filters [58] address this difficulty by guaranteeing successful construction while retaining near-optimal space usage, making them a promising foundation for compact interleaved indexes at scale. The contribution of this paper is threefold. First, we introduce an approximate colored index based on interleaved ZOR filters, combining guaranteed filter construction with near-optimal filter space. Second, we introduce a color-preserving merging mechanism that extends fingerprints with membership bits, allowing multiple genomes to share a filter while retaining their individual identities. Together with a lightweight, sketch-based layout planner, this mechanism exploits sequence sharing while accommodating collections with widely varying genome sizes and degrees of similarity. Third, we support low-density minimizer sampling alongside gapped *k*-mers and randstrobes, allowing users to tailor indexing costs and sensitivity to their data and application characteristics. We implement these advances in Zindex, a fast, parallel, SIMD-accelerated Rust index that scales to millions of genomes while requiring low memory.

## 2 Methods

A filter is a probabilistic data structure that compactly represents a set *S* and answers membership queries, always reporting elements of *S* as present but occasionally also reporting an element outside *S* as present, an error called a *false positive*. This query semantics is called *approximate membership query* (AMQ), and the false positive rate (FPR) is a user-chosen parameter *ϵ* [36,59]. A space-efficient way of guaranteeing this FPR is to store *f*-bit fingerprints, where *f* = ⌈–log_2_ (*ϵ*)⌉, in an array *A* of |*A*| = ⌈*c*· |*S*|⌉ slots, using a small number of hash functions to guide insertions and probes, e.g., three: *H* = (*h*_1_, *h*_2_, *h*_3_) [60,55,56,58]. The array requires |*A*| · *f* ≈ *c* · |*S*| · *f* bits in total, where the size factor *c* approaches the information-theoretic lower bound of 1 in state-of-the-art filters [61,58,62].

AMQ semantics provides a modular framework for approximate sequence search, exemplified by threshold queries that report a target genome whenever the fraction of shared query *k*-mers reaches a specified cutoff *θ* [37]. Filters are agnostic to the underlying sequence sampling and seeding scheme, so either can be changed without changing the underlying data structure (Section 2.3). In contrast, exact methods based on unitigs or spectrum-preserving string sets compress *k*-mers by exploiting the overlap between consecutive ones, an advantage that sampled keys forgo [29,31,30].

Consider an AMQ-based colored index for *N* documents, with one filter per document sized to its number of distinct *k*-mers (or minimizers). For a query containing *x* keys, searching all documents requires *xN* filter lookups, each accessing up to |*H*| array positions. If every access misses the cache, this produces up to *xN* |*H*| cache misses per query, making memory access a bottleneck for large collections. Several complementary strategies can reduce this cost: improving the memory layout of filters (Section 2.1), organizing documents into fewer filters while retaining document-level results (Section 2.2), and sampling fewer keys (Section 2.3). Together, these design choices motivate Zindex, whose architecture is summarized in Figure 2.

### 2.1 Interleaved ZOR Filter

Interleaved filters store slot *i* of every filter contiguously, so that the *N* slots probed by a hash function lie next to each other in memory (Figure 1a), yielding two benefits. First, each hash probe reads a single row of *N f* bits, touching only ⌈*N f* / *B*⌉ cache lines instead of *N*, where *B* is the cache line size in bits [42,41]. Second, the probed rows can be processed with SIMD: for each key, the |*H*| probed rows are XORed and compared against the query fingerprint in a single fused pass, without materializing an intermediate row. With AVX2, a single 256-bit operation screens 32 filters with 8-bit fingerprints. This layout imposes two requirements: (i) *h*_*i*_ should be the same hash function for every filter; and (ii) *h*_*i*_ (*x*) mod |*A*| should give the same array index, meaning that every filter array must have the same size.

**Fig. 1:**
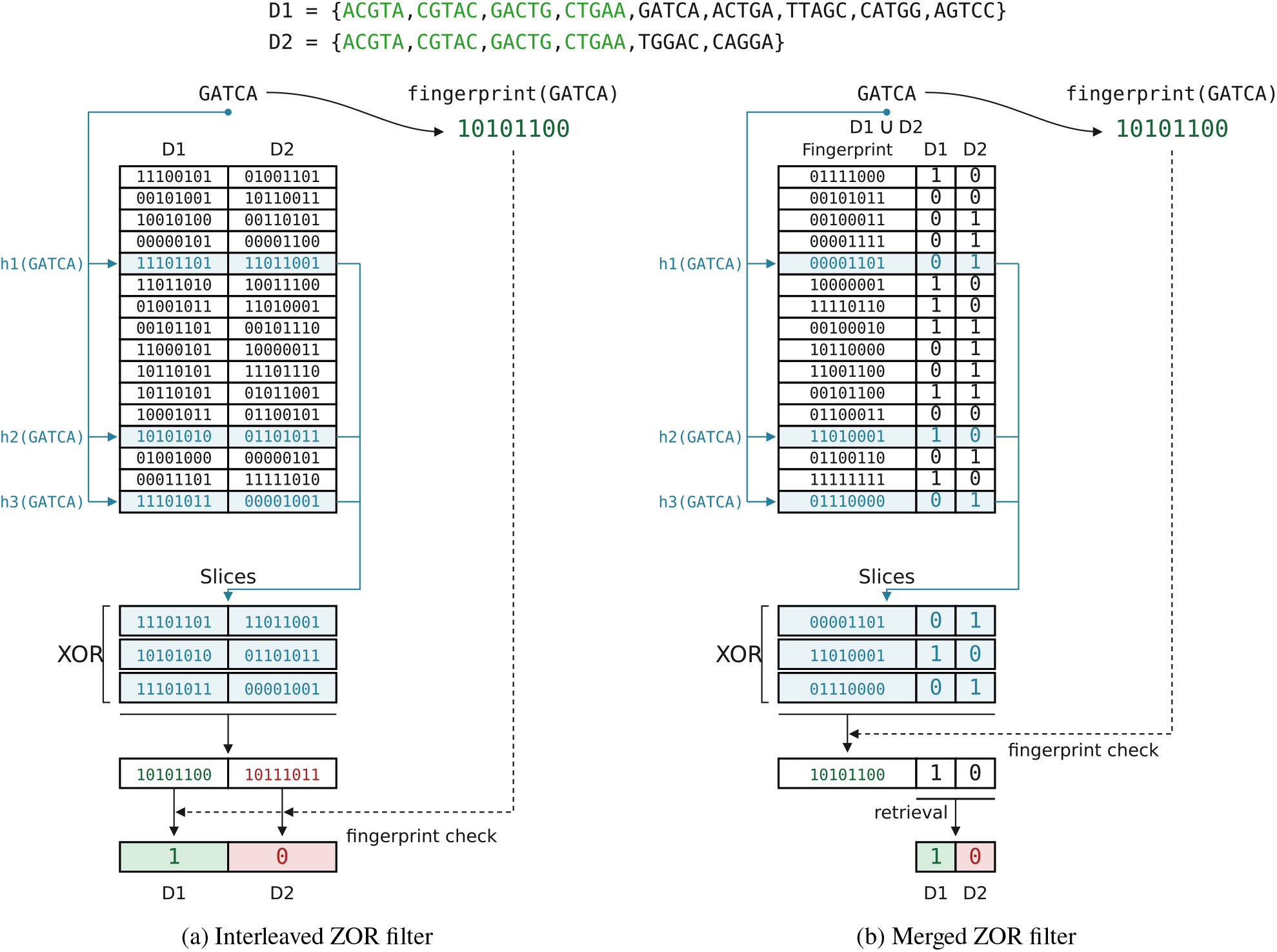
Power-of-two sized ZOR filters with 8-bit fingerprints. (a) Interleaved filter. The largest document, *D*_1_ with |*D*_1_| = 9, sets the size to 16 slots, for a total of 2 × 16 × 8 = 256 bits. A query XORs the three slots selected by *h*_1_, *h*_2_, *h*_3_ and compares each document’s resulting fingerprint with that of the query item, one comparison per document. (b) Merged filter. |*D*_1_ ∪ *D*_2_| = 11, so the size remains 16 slots, for a total of 16×(8+2) = 160 bits, a 37.5% saving. A query XORs the same three slots, but performs a single fingerprint comparison; on a match, the resulting bitmask gives the documents containing the item.

Binary fuse [56] and XOR filters [55] are XOR-based, highly space-efficient, static alternatives to Bloom filters [36]. Binary fuse filters can even be as small as 75% of a Bloom filter with matched settings. Moreover, they do not require additional hash functions to achieve a lower false-positive rate, unlike Bloom filters, further motivating their use.

XOR-based filters share the following query procedure: There is a small, constant number of hash functions, e.g., three: *H* = (*h*_1_, *h*_2_, *h*_3_) ; a fingerprint function *F* that outputs a *f*-bit fingerprint; and an array *A*, where each slot stores a partial fingerprint. For a query item *x*, the query returns “found” only if the complete stored fingerprint *A*[*h*_1_ (*x*)] ⊕ *A* [*h*_2_ (*x*)] ⊕ *A* [ *h*_3_ (*x*)] equals the query fingerprint *F(x)*. Figure 1a shows this query on an interleaved filter over two documents: for `GATCA,` the three probed rows XOR to *F* (`GATCA`) = `10101100` in *D*_1_’s column, but not in *D*_2_’s. An item that is not in *S* incorrectly returns “found”, i.e. a false positive, with a probability of only 2^– *f*^ due to the random nature of the fingerprint function. Binary fuse filters partition *A* into segments of power-of-two size and map each key to slots in consecutive segments, improving locality and allowing higher occupancy than conventional XOR filters [56]. In these structures, increasing the number of hashes can reduce |*A*| without changing the false-positive rate [63,64], while increasing the fingerprint length *f* lowers the false-positive rate without requiring additional hashes.

XOR and binary fuse filters use peeling-based construction algorithms [55,56] that assign values to *A* so that the query equations hold for every stored key. For a given key set *S*, construction succeeds with high probability but may fail for the selected hash functions *H*, requiring a new choice *H*^′^ and another construction attempt. Such failures are manageable for independent filters, since only the failed filter needs to be rebuilt, but interleaving requires a shared *H* that succeeds for all *N* filters. A failure in any filter can therefore force previously completed filters to be rebuilt under new hash functions, making even rare individual failures costly when many filters are interleaved. Taxor [57] successfully interleaves XOR filters but identifies these repeated builds as an obstacle to adopting the more space-efficient binary fuse filters, whose higher construction failure rates amplify this problem.

Rather than selecting new hash functions when peeling stalls, ZOR filters [58] set aside selected keys and continue construction under the same *H*, preserving completed work for the current filter (and the other interleaved filters in our case). The main array therefore represents a subset *S*^′^ ⊂ *S*, while an auxiliary structure stores the abandoned keys to preserve membership guarantees for the full set *S*. This guarantees successful construction and allows the main array to contain just one slot per original key, | *A*| = | *S*|, with only a small fraction of keys requiring auxiliary storage. Although a single ZOR filter queries its auxiliary structure only when its main filter returns negative, querying *N* independent filters can still require up to *N* auxiliary lookups per key, since a key is rarely present in all *N* documents. To avoid this overhead, we interleave the main arrays while replacing the individual auxiliary structures with a single global retrieval structure that maps each abandoned key to the list of filters that abandoned it. For each query key, Zindex unconditionally queries this global structure once alongside the interleaved filters, replacing up to *N* auxiliary membership checks with a single lookup. In Zindex, this global structure is a hashmap of 32-bit key hashes, which increases the per-document FPR negligibly while trading a small amount of space for query speed. Although ZOR reduces the space needed to represent each key, interleaving still requires equally sized arrays, so filters representing smaller sets inherit the capacity required by the largest set. Avoiding this waste on collections with widely varying dataset sizes requires a layout that balances the number of distinct keys represented by each filter within an interleaved group.

### 2.2 Layout

Hierarchical interleaved filters [57,47,45] accommodate differences in document cardinality by splitting large documents into fragments and grouping smaller ones within a hierarchy. In hierarchical layouts, recovering the individual sources of a hit to a merged filter requires descending into additional filters, increasing storage, construction work, and query cost. Splitting a document across several filters can also increase its false-positive probability when each query key must be tested against all fragments. For *s* fragments with individual false-positive probability *ϵ*, the combined probability is bounded by *sϵ* and equals 1 – (1 –*ϵ*)^*s*^ under independence. Maintaining the original bound therefore requires more selective fragment filters, with ⌈log_2_ *s*⌉ additional fingerprint bits sufficient for fingerprint-based filters.

To preserve document identities directly, we extend ZOR filters with associated values, following the principle of static retrieval structures [65,61]. For a cluster of *n* > 1 documents, each indexed key is associated with its *f*-bit fingerprint and an *n*-bit membership mask identifying the documents that contain it. Construction assigns these combined values to the filter slots, so that XORing the probed slots retrieves both the fingerprint and the membership mask. Each distinct key in the union is represented once, while its mask retains all its document memberships. Singleton filters omit the membership mask, giving *r* (1) = *f* and *r* (*n*)= *f* + *n* for *n* > 1. We require *n* ≤ 64 – *f* so that each logical slot value fits within a 64-bit word. For a key absent from the cluster, a false hit to a particular document requires both a matching fingerprint and a set membership bit. The main-filter false-positive probability per document is therefore at most 2^–*f*^.

Whether merging saves space depends on both sequence sharing and the additional membership bits. We estimate sharing using sketches adapted from SimdSketch [66] to operate on indexed feature IDs and support unions. Jaccard similarity is estimated from the fraction of matching fingerprints, while a merged sketch is updated by taking the component-wise minima of its members’ sketches. For two clusters with *n*_1_ and *n*_2_ documents, current union-cardinality estimates *c*_1_ and *c*_2_, and estimated Jaccard similarity 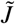, we estimate their combined union cardinality as 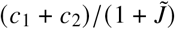. We clamp this estimate to [max(*c*_1_, *c*_2_), *c*_1_ + *c*_2_] and denote the result by Ĉ. Ignoring capacity rounding, merging is beneficial when

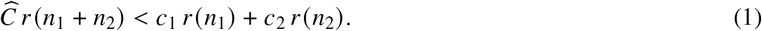

Thus, high similarity alone does not justify a merge, because the savings from shared keys must exceed the cost of widening the slots.

We choose to round capacities to powers of two to enable fast modulo reduction by bit masking and group filters into a small number of interleaved subindexes, trading padding for fewer row probes. The implemented criterion also accounts for the power-of-two capacities used for interleaving, defining *P*(*x*) as the smallest power-of-two capacity that is at least *x*:

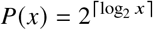

If the two clusters currently have planned capacities *m*_1_ and *m*_2_, a merge is accepted only when

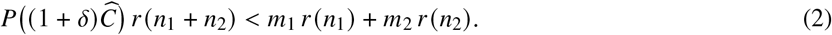

The planning margin *δ*, set to 0.03 by default, provides headroom against underestimating a union near a power-of-two boundary (See the *δ* = 0 ablation in Supplementary S8).

We implement two greedy planners using this same cost model. The full planner initially evaluates all document pairs and repeatedly selects the admissible merge with the largest estimated space saving. After each merge, it evaluates candidates involving the new cluster while retaining previously computed candidates between unchanged clusters. This provides a straightforward planner for smaller collections, but requires *O* (*N*^2^) sketch comparisons and potentially *O* (*N*^2^) storage for merge candidates. For larger collections, the default streaming planner processes documents in input order, merging each incoming document into the existing cluster that offers the largest positive estimated saving or creating a singleton cluster when no merge is beneficial. To avoid comparing every incoming document with every cluster, it groups existing clusters by filter capacity and document count, which together determine their storage cost and the slot width required after merging. An optimistic bound on the possible saving then excludes entire groups whose candidates cannot improve the current best merge, without computing their similarities. Because it preserves input order, labeled collections can be streamed in taxonomic order to bring related documents close together, following the motivation of phylogenetic compression [67] (See the input order ablation in Supplementary S8).

Once clustering is complete, we construct the exact unions and membership masks and determine filter capacities from their actual cardinalities. The resulting singleton and merged filters are grouped by their power-of-two capacities, allowing filters with different slot widths to share an interleaved subindex (Figure 2). The clustering postprocessing can enforce a minimum number of documents per class and a cap on the number of classes to trade additional space for fewer subindex probes. Within each subindex, every row begins with the contiguous *f*-bit fingerprints of all filters, followed by the membership fields of merged filters. Queries XOR and compare the fingerprint regions using SIMD, retrieving and XORing membership fields only for filters whose fingerprints match. Within equal-width groups, columns are ordered using their retained union-sketch fingerprints to improve compression.

**Fig. 2:**
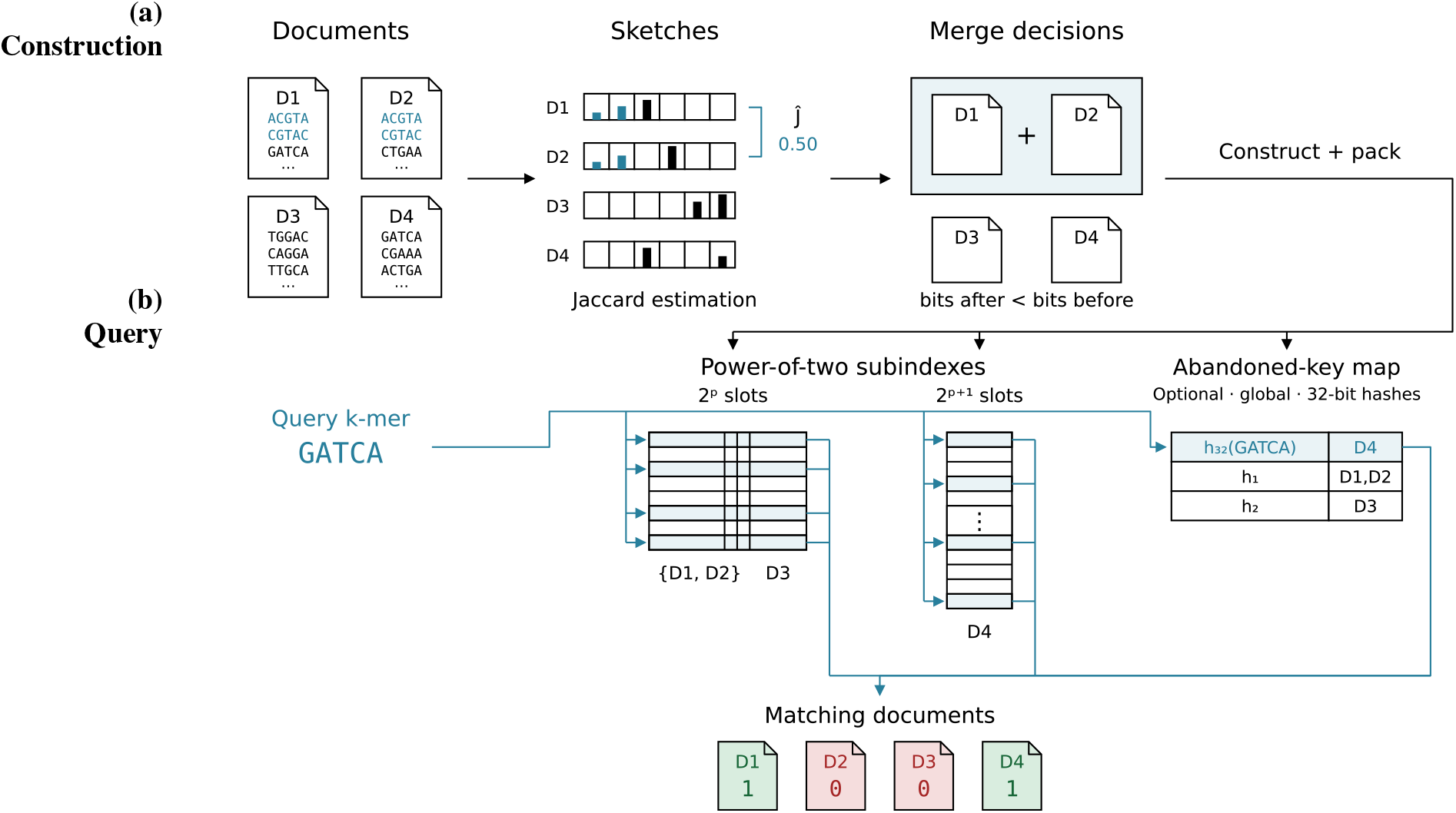
Overview of Zindex. (a) Construction. Documents are sketched, and similar ones are merged into clusters (*D*_1_ and *D*_2_); the resulting filters are partitioned into power-of-two size buckets (subindexes), and abandoned keys are stored in a global map.

To bound the payload of individual subindexes, oversized size classes are further partitioned by a deterministic hash of the indexed key. This preserves the document clusters while routing each key to exactly one partition within its size class, so partitioning does not multiply the number of membership tests for a document. A document’s keys may therefore occupy several physical partitions, but each query key probes only its assigned partition.

Subindexes are stored as independently decompressible zstd-compressed frames, together with the shared abandoned-key recovery structure described above. By default, queries process one subindex at a time and combine its hits with auxiliary recovery results, limiting resident filter memory while accumulating matches for each document. A document is reported when its matching fraction reaches the threshold *θ* over the extracted query keys.

### 2.3 Seeding and sampling schemes

In sequence indexing, *seeding* determines the construction of the extracted features, such as contiguous *k*-mers, spaced seeds, or strobemers, whereas *sampling* selects which features are retained, using schemes such as minimizers or FracMinHash. Because these operations ultimately produce hashable keys, AMQ filters can accommodate different combinations of seeding and sampling without changing the index structure. Despite this, few AMQ-based tools exploit sampling and alternative seeds: the SeqAn-based Raptor [50] supports minimizer sampling and gapped *k*-mers, while the taxonomic classifier Taxor [57] supports minimizer and syncmer sampling. In contrast, many others, such as the sequence Bloom tree family [37,40], COBS [46], and kmindex [48] index all *k*-mers. A random minimizer scheme [32] selects the smallest *k*-mer under a fixed random hash order from each window of *w* consecutive *k*-mers, giving an expected sampling density of approximately *d* = 2/ (*w*+ 1). Modern low-density schemes [51,52] can achieve densities closer to the (*k, w*)-dependent lower bounds established for forward sampling schemes [68]. Applying the same deterministic selector to references and queries guarantees that every exact shared region of length at least *k*+*w*–1 contains a shared minimizer, but does not guarantee that a partially matching query passes a sequence-level threshold. Zindex supports random minimizers [32] computed with SimdMinimizers [69], open-closed mod-minimizers [52], and hash-based sampling as in FracMinHash [70]. Both minimizer selectors have SIMD implementations, while SIMD-accelerated input parsing [71] is shared across extraction modes. For dense and minimizer indexing, Zindex additionally supports symmetric gapped *k*-mers masks [53] and randstrobemers [54], with minimizer selection performed before masking or linking when sampling is enabled.

## 3 Results

### 3.1 Experimental setup

Experiments used four machines: M1 with 64 GB of RAM and a 2 TB SSD, M2 with two Xeon Gold 6330N CPUs (56 cores), 811 GB of RAM and two 7200 rpm SATA HDDs of 8 TB and 6 TB, M3 with four Xeon E7-4830 CPUs (32 cores, without AVX2), 540 GB of RAM and a 6 TB, 7200 rpm SATA HDD, and M4 with Intel Xeon Gold 6430 CPUs (32 cores), 540 GB of RAM and SSD storage. Every comparative experiment was limited to 24 hours, builds exceeding this limit were discarded and not queried.

We compared Zindex with four approximate colored indexes: COBS [46], based on interleaved Bloom filters grouped into size-matched blocks; Raptor [50,47], based on hierarchical interleaved Bloom filters; kmindex [48], based on a partitioned Bloom-filter matrix; and Kaminari [72], which represents minimizer color sets with fingerprint-based membership checks. We complemented this comparison with three exact colored indexes: Fulgor [31], which maps unitigs to compressed color sets; Themisto [73], based on the spectral Burrows–Wheeler transform with compressed color sets; and MetaGraph [11], which combines the BOSS succinct de Bruijn graph [74] with compressed annotations.

We first compared the tools on nested samples from RefSeq [75] and AllTheBacteria (ATB) [76], beginning with 64 genomes and extending to larger collections for tools that remained feasible. The RefSeq selection emphasizes taxonomic diversity, whereas the ATB samples retain related assemblies, allowing us to examine performance on collections with different degrees of redundancy. These experiments used 32 threads on M1, compressed inputs, *k* = 31 and a target per-key FPR of 2^–8^ unless stated otherwise, with probabilistic settings calibrated on a standalone 5 Mb genome and then fixed throughout collection growth. We compared full *k*-mer indexing with minimizer indexing, favoring native minimizer implementations where available; for COBS, kmindex, Fulgor, MetaGraph and Themisto, we precomputed random minimizers with SimdMinimizers [69] and supplied the resulting FASTA files. Second, the tools whose minimizer indexes scaled up to 16k genomes within the limit, Raptor and COBS, were run against Zindex on two complete collections: the 661,405 bacterial assemblies of the 661k collection [77] and the 268,827 *Mycobacterium tuberculosis* contig sets assembled by Logan [12]. These runs used 32 threads on M2 and minimizers of 19-mers from 31-base windows. Finally, Zindex alone indexed the eukaryotic subset of RefSeq, the full RefSeq, and all 2,763,297 genomes of ATB.

Positive and negative workloads each comprise 16,384 sequences of 1 kbp, with positive fragments sampled from indexed genomes and synthetic negatives generated as uniformly random DNA. Dense indexes were queried with all *k*-mers, with native minimizer computation where available and precomputed minimizers otherwise, with an 80% matching threshold throughout. Query times on sampled collections are medians of three runs, while complete-collection measurements use cold starts after eviction of the index from the page cache.

### 3.2 Comparison on sampled collections

#### Index size and construction

Minimizer indexing substantially reduces the size of the approximate indexes, while not reliably benefiting the exact ones: supplying minimizer FASTA files increases the size of both Fulgor and Themisto, and MetaGraph changes only modestly (Fig. 3; for clarity, only their dense indexes are shown). Filters pay a roughly fixed cost per key, so their size falls with the number of keys, whereas exact indexes compress by exploiting the overlap between consecutive *k*-mers, which sparse minimizers rarely share (Section 2). For Zindex, sampling reduces index size by more than an order of magnitude relative to dense indexing, yielding the smallest minimizer index at every shared collection size in this RefSeq comparison. Among the minimizer indexes constructed for 16,384 genomes, Zindex was also the fastest to build, ahead of COBS and Raptor (Fig. 4a). Memory-related failures stopped larger builds of MetaGraph, Kaminari, Raptor, Fulgor and dense Zindex, whereas disk-related issues limited Themisto and kmindex.

**Fig. 3:**
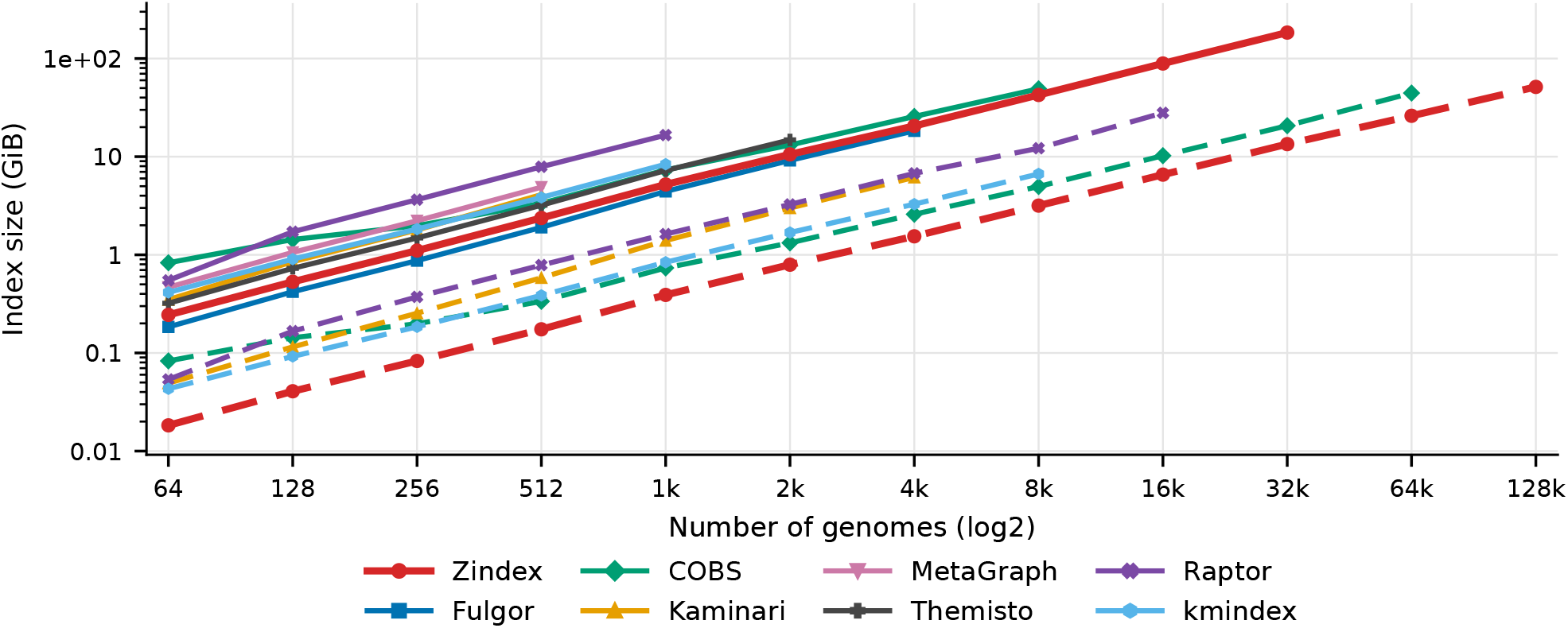
Index size on diverse RefSeq collections at target FPR 2^–8^. Solid lines denote *k*-mer indexes and dashed lines minimizer indexes. Minimizers select 31-mers from 49-base windows.

**Fig. 4:**
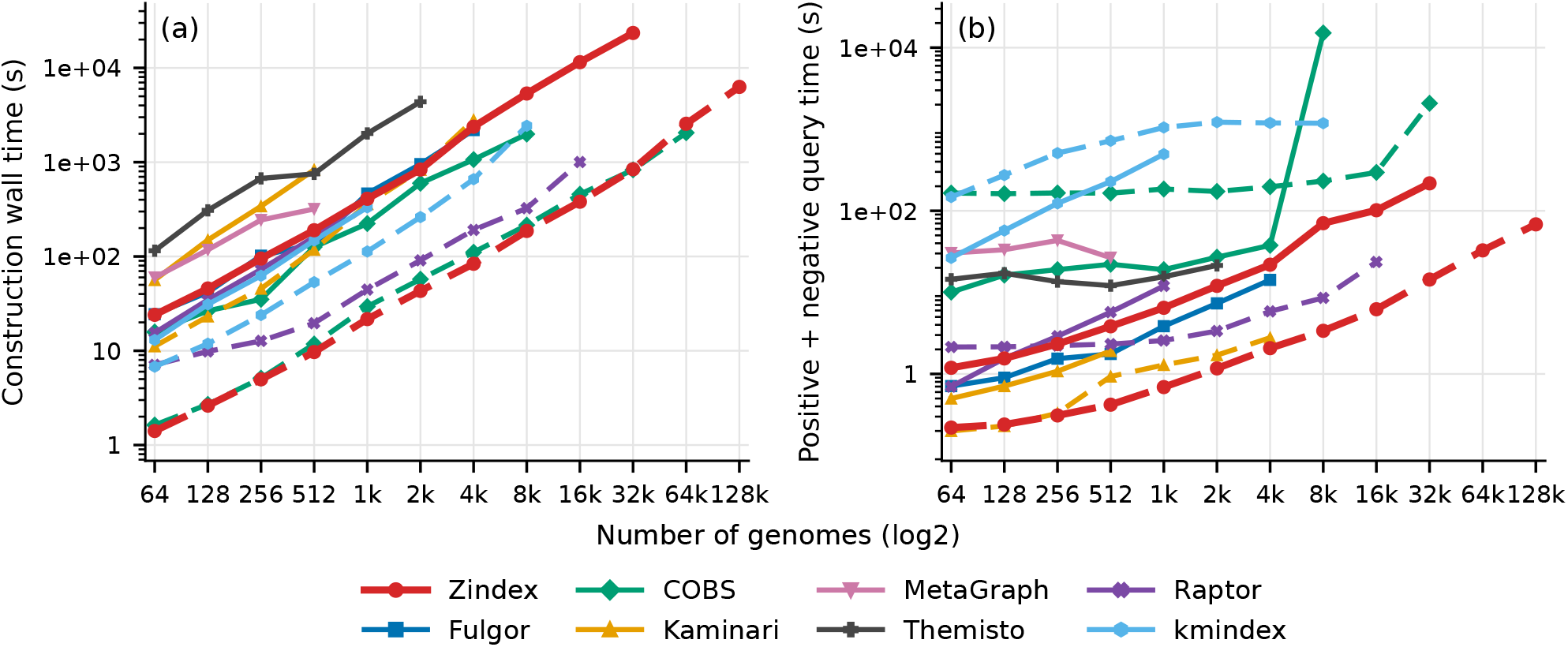
RefSeq performance at target FPR 2^–8^: (a) construction wall time and (b) combined positive and negative query wall time. Query time is the wall time of the positive workload plus that of the negative workload, each the median of three runs including index loading and output. Solid lines denote dense indexing and dashed lines minimizer indexing. External minimizer preparation is excluded, while native minimizer computation is included.

#### Query performance

Figure 4b reports the combined positive and negative query wall time; separate results are given in the Supplementary Results. Zindex achieves the lowest combined minimizer query time across all shared collection sizes except 4,096 genomes, where Kaminari is slightly faster. At the largest collection, Zindex queries approximately twice as fast as Raptor and more than an order of magnitude faster than COBS, and its positive and negative query times remain similar throughout collection growth (Supplementary Fig. S2). In dense mode, Fulgor queries approximately twice as fast as Zindex on the larger shared collections, and Kaminari approximately twice as fast on the smaller collections for which its dense index completed; Themisto and MetaGraph are slower than Zindex, and kmindex is substantially slower in both modes. The abrupt slowdown of dense COBS at 8,192 genomes occurs when its index exceeds the available RAM, so that query processing becomes dominated by disk I/O. Its smaller minimizer index still fits in memory and avoids this slowdown, although its query times remain substantially higher than those of Zindex (Supplementary Fig. S2). Overall, Zindex provides the fastest minimizer queries at the largest tested scales, while Fulgor and Kaminari retain an advantage when querying dense indexes.

#### Collection redundancy and precision

We next examined how collection redundancy and the target false-positive rate affect the relative performance of the indexes. On the redundant ATB samples, Fulgor’s dense-space advantage over Zindex increases, while Themisto and Kaminari also become more competitive in space (Supplementary Fig. S4). Nevertheless, at the largest 8-bit ATB sample, Zindex produces the smallest minimizer index and achieves the lowest construction and combined query wall times among the tools that completed the benchmark. Relaxing the target FPR from 2^–8^ to 2^–4^ reverses the dense-index comparison on RefSeq, with Zindex becoming smaller than Fulgor (Supplementary Figs. S1 and S2). Thus, when a higher per-key false-positive rate is acceptable, approximate indexing can also provide a space advantage without sampling. At the largest completed RefSeq samples, Zindex also provides the smallest minimizer index and the lowest construction and combined query wall times at both 4-bit and 16-bit precision (Supplementary Figs. S1 and S3). At 16-bit precision on ATB, the comparison with Kaminari is closer: at the largest available sample, Kaminari produces a slightly smaller minimizer index and answers queries faster, whereas Zindex constructs its index substantially GB are decimal; RAM is peak process RSS. Runs used 32 threads on M2 unless marked: ^*a*^ 64 threads on M4; ^*b*^ 32 threads on M3, not subject to the 24-hour limit. OCM denotes open-closed mod-minimizers and Random denotes random minimizers. Raptor was run with union estimation and rearrangement disabled.

faster (Supplementary Fig. S5). Against COBS and Raptor, Zindex retains a clear advantage on ATB at this precision, with a smaller minimizer index and lower construction and query wall times. Redundancy therefore makes Kaminari a stronger competitor at high precision without removing Zindex’s advantages over the Bloom-filter-based indexes.

### 3.3 Seed robustness and computational cost

To assess how seed choice affects robustness to sequence errors, we compared query-base coverage for contiguous 21-mers, a spaced seed selecting 21 of 25 positions, and randstrobemers joining two 11-base strobes, with and without minimizer sampling. Queries were generated from paired 10 kbp parent reads from 32 genomes under substitution, insertion, deletion and mixed error profiles.

As expected, spaced seeds improve coverage under substitutions, whereas randstrobemers retain greater coverage under insertions and deletions, with this advantage extending to the mixed error profiles (Fig. 5a–e). Minimizer sampling retains the qualitative differences between seed types while reducing index size and construction time by roughly an order of magnitude (Fig. 5f,g). The three dense seed types have similar storage and construction costs. Query cost exposes a further trade-off: spaced seeds remain close to contiguous seeds, whereas randstrobe queries are approximately twice as expensive (Fig. 5h), because each randstrobe requires a second strobe to be selected among candidate positions before hashing. These results show that the same index structure can accommodate seeds tailored to different error profiles without substantial storage or construction overhead.

**Fig. 5:**
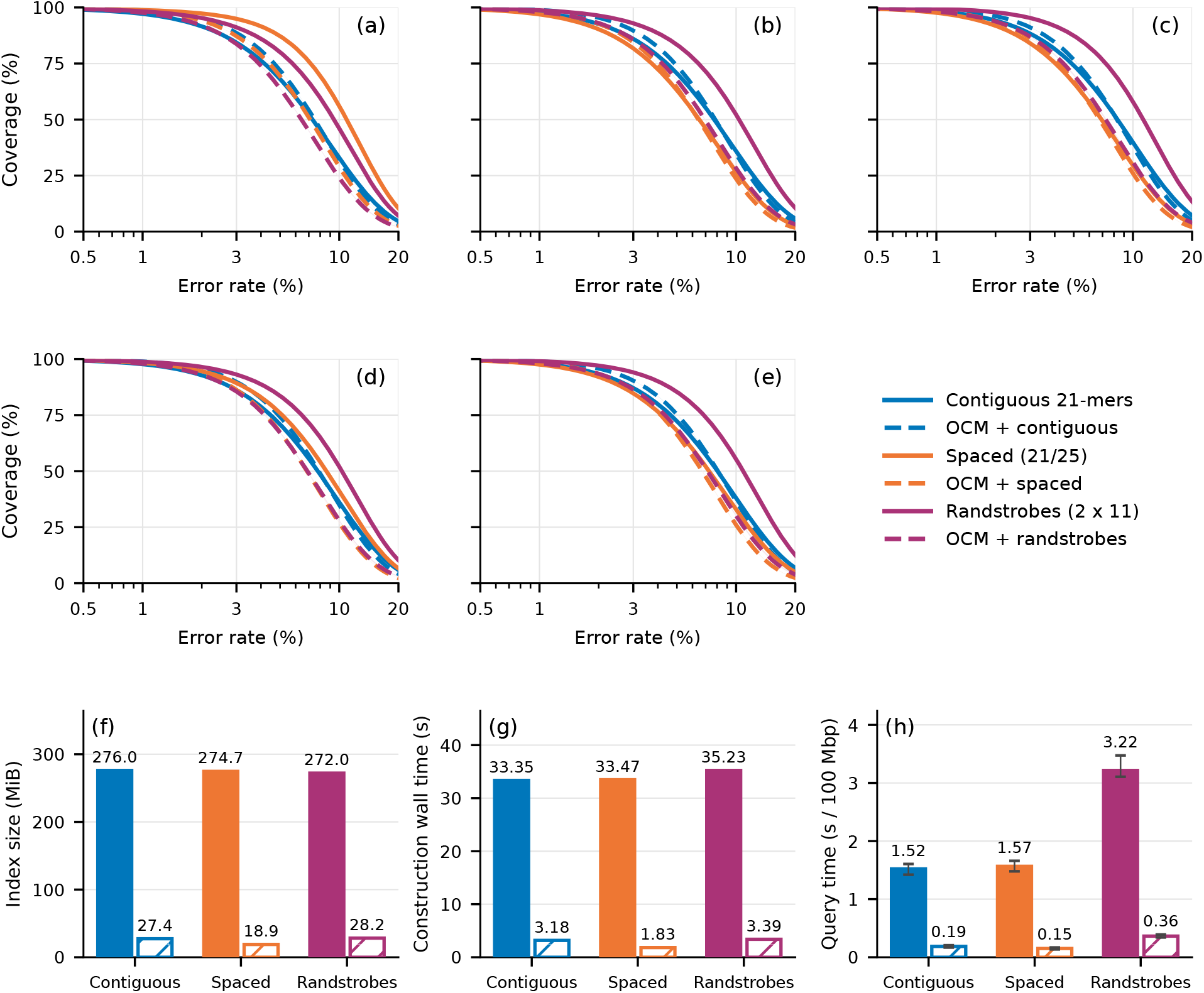
Seed robustness and cost on 32 genomes. Approximate query-base coverage under (a) substitutions, (b) insertions, (c) deletions, (d) ONT-like and (e) HiFi-like mixtures. Bands are 95% bootstrap intervals over 16 genome pairs. Queries use 3,200 paired 10 Kbp parent reads. Coverage is measured over query bases, with sampled matches marking their represented windows and dense matches marking the positions selected by matching seeds. The simplified ONT-like and HiFi-like profiles use substitution:insertion:deletion ratios 39:24:36 and 2:10:13, respectively. Panels (f)–(h) show index size, construction wall time, and query time with 32 threads. Hatched bars use minimizers. Timings exclude parsing, and whiskers show min/max envelopes over paired conditions and repetitions.

### 3.4 Scaling to complete collections

#### Comparison on 661k and M. tuberculosis

Table 1 summarizes construction on the two complete comparative collections. On 661k, no Raptor configuration completed: the default configuration exhausted the 811 GB of M2 after 2.8 hours, and the configuration without union estimation and rearrangement was discarded at 24 hours. Zindex built the 661k index in 11.4 hours with a peak of 33.1 GB of memory, producing a 97.0 GB index. On the redundant *M. tuberculosis* collection, Raptor completed only with union estimation and rearrangement disabled. At the matched per-key FPR of 2^–8^, Zindex’s index was 5.5 times smaller than Raptor’s (50.5 versus 277.3 GB) and was built 9.5 times faster (1.1 versus 10.3 hours) with 8 times less memory (35.3 versus 283.3 GB). Even at its default FPR of 0.05, about 13 times higher, Raptor produced a 178.0 GB index, 3.5 times larger than Zindex’s. To distinguish the contribution of the index structure from that of sampling, we repeated the build with random minimizers, obtaining a 71.7 GB index that remained 3.9 times smaller than Raptor’s at the same FPR. Open-closed mod-minimizers provided a further 30% reduction in index size and reduced construction time from 2.6 to 1.1 hours. COBS with minimizer extraction exceeded 24 hours on this collection at its default FPR of 0.3. Supplementary Table S1 lists all runs, including Zindex ablations of input order, hash count, merge margin and thread count. At this scale, loading dominates query wall time: both tools read their index from disk at about 180 MB/s, so load time is proportional to index size (Table 2). On *M. tuberculosis*, Zindex consequently answered the combined workload 3.5 times faster than Raptor at the same FPR (15.2 versus 53.4 minutes) with at most 105.9 GB of memory, against 277.6 GB for Raptor. Although its larger index increased cold-start time, Raptor processed the positive workload faster after loading. Under linear extrapolation of the reported loading and search times, its total time would match Zindex’s at approximately 64,000 positive reads. Positive reads in this collection match a large fraction of the 268,827 indexed runs, and to limit memory Zindex loads one subindex at a time, keeping partial per-read answers until all subindexes are processed. The higher wall time and memory of the positive workload come from the answers themselves, which cover most of the 268,827 runs for every read, whereas negative reads are rejected quickly by both tools. On 661k, Zindex answered each workload in about ten minutes, over 90% of which was spent loading the index.

**Table 1:** Construction on the complete collections with minimizers of 19-mers from 31-base windows, four hash locations and 8-bit fingerprints.

| Collection | Assemblies | Tool | FPR | Index (GB) | Wall (h) | RAM (GB) |
|---|---|---|---|---|---|---|
| 661k | 661,405 | Zindex (OCM) | 2<sup>−8</sup> | 97.0 | 11.4 | 33.1 |
| *M. tuberculosis* | 268,827 | Zindex (OCM) | 2<sup>−8</sup> | 50.5 | 1.1 | 35.3 |
|  |  | Zindex (Random) | 2<sup>−8</sup> | 71.7 | 2.6 | 65.0 |
|  |  | Raptor | 2<sup>−8</sup> | 277.3 | 10.3 | 283.3 |
|  |  | Raptor | 0.05 | 178.0 | 11.0 | 185.5 |
| Eukaryotic RefSeq<sup>*a*</sup> | 2,439 | Zindex (OCM) | 2<sup>−8</sup> | 178.5 | 10.0 | 394.2 |
| Full RefSeq<sup>*a*</sup> | 564,401 | Zindex (OCM) | 2<sup>−8</sup> | 289.5 | 21.7 | 399.2 |
| Full ATB<sup>*b*</sup> | 2,763,297 | Zindex (OCM) | 2<sup>−8</sup> | 406.3 | 95.5 | 43.8 |

GB are decimal; RAM is peak process RSS. Runs used 32 threads on M2 unless marked: <sup>*a*</sup>64 threads on M4; <sup>*b*</sup>32 threads on M3, not subject to the 24-hour limit. OCM denotes open-closed mod-minimizers and Random denotes random minimizers. Raptor was run with union estimation and rearrangement disabled.

**Table 2:** Cold-start query wall time and peak memory on the complete collections for 16,384 sequences of 1 Kbp at an 80% threshold, on M2 with 32 threads.

| Collection | Tool | Workload | Wall (min) | Load (s) | Search (s) | RAM (GB) |
|---|---|---|---|---|---|---|
| 661k | Zindex | positive | 10.8 | 546 | 101 | 90.3 |
|  |  | negative | 9.2 | 545 | 8 | 80.5 |
| *M. tuberculosis* | Zindex | positive | 10.5 | 279 | 350 | 105.9 |
|  |  | negative | 4.7 | 279 | 2 | 34.2 |
|  | Raptor (2<sup>−8</sup>) | positive | 26.9 | 1,587 | 17 | 277.6 |
|  |  | negative | 26.6 | 1,587 | 2 | 277.4 |
| ATB | Zindex | positive | 55.6 | 2,750 | 581 | 418.9 |
|  |  | negative | 45.4 | 2,690 | 31 | 378.1 |

Wall time includes index loading and output; load and search are each tool’s internal timers. The ATB index was built on M3 and queried on M2.

#### Repository-scale constructio

. We then evaluated Zindex alone on complete repositories, extending the bacterial experiments to the heterogeneous genome sizes represented in RefSeq (Table 1, bottom). The complete RefSeq snapshot dated 20 October 2026, comprising 564,401 assemblies, was indexed in 21.7 hours, producing a 289.5 GB index. The complete ATB collection extends them to millions of documents: Zindex indexed all 2,763,297 genomes in 95.5 hours with a peak of 43.8 GB of memory, producing a 406.3 GB index. This build ran on M3 without AVX2 and was not subject to the 24-hour limit, so its running time is an upper bound for SIMD-capable hardware. The layout planner sketched, ordered and planned the whole collection in 26.2 hours within the same memory budget, even though the final index occupies 690 GB before compression. Querying the ATB index on M2 required 378–419 GB of memory and 45–56 minutes per workload, of which 45 minutes was spent loading (Table 2). These results demonstrate repository-scale construction with modest memory, while the measured query configuration still incurs substantial loading and memory costs, motivating the query optimizations discussed in Section 4.

## 4 Future work and conclusions

We have introduced interleaved ZOR filters as a foundation for approximate colored indexing, combining guaranteed construction with near-optimal filter space. Building on this structure, our color-preserving merging mechanism retains individual dataset identities while exploiting sequence sharing, and our sketch-based layout algorithm accommodates collections with widely varying cardinalities and similarities. The resulting hash-based framework supports low-density sampling and sensitive seeds without structural changes, allowing the indexed features to be adapted to different data and applications. Zindex brings this framework to repository scale, combining parallelism and SIMD in Rust to deliver feasible construction across millions of genomes with very low memory requirements. Across the evaluated collections, Zindex combined compact minimizer indexes with fast construction, scaling from diverse bacterial samples to complete public genome collections. On 268,827 *M. tuberculosis* assemblies, it produced an index 5.5 times smaller than Raptor’s at the same false-positive rate, with 9.5 times faster construction and eightfold lower peak memory. At repository scale, Zindex indexed the tested RefSeq snapshot in under 22 hours and all 2,763,297 genomes of AllTheBacteria in four days using 43.8 GB of peak construction memory. With construction now practical at repository scale, reducing query memory traffic becomes the next priority for improving performance without sacrificing filter space efficiency. The current lookup combines |*H*| *f* fingerprint bits per filter, raising the question of whether fewer bits or memory blocks can be accessed with comparable space efficiency and retaining efficient SIMD processing. A complementary approach, following PAC [49], would augment the index with a compact auxiliary structure to rule out irrelevant clusters or subindexes, accelerating queries by avoiding unnecessary filter probes at a small additional space cost.

Alongside these query optimizations, further space gains require exploiting redundancy across the collection, beyond the efficiency of individual filters. Our merging mechanism shares keys within clusters, but keys and membership patterns can still be repeated across clusters, motivating representations that share fingerprints or encode repeated color sets [72,31]. The central question is therefore whether layouts built from document filters can capture this remaining redundancy, or whether a more global representation of the collection is needed.

### LLM Usage Statement

Claude Opus, Fable(Anthropic), Codex, and Gemini Pro were used to assist with the implementation of Zindex and to rephrase sentences and correct grammar in the manuscript. All generated code and text are inspected, reviewed, and corrected by the authors.

