## Supplementary Material for "Toward space-optimal approximate colored indexes at scale"

### S1 RefSeq at 4-bit precision

At lower precision, Zindex's dense index becomes more competitive with Fulgor, while its minimizer index remains compact relative to COBS, Raptor and Kaminari. COBS often uses less construction memory; Zindex combines short sampled construction with much faster batch queries.

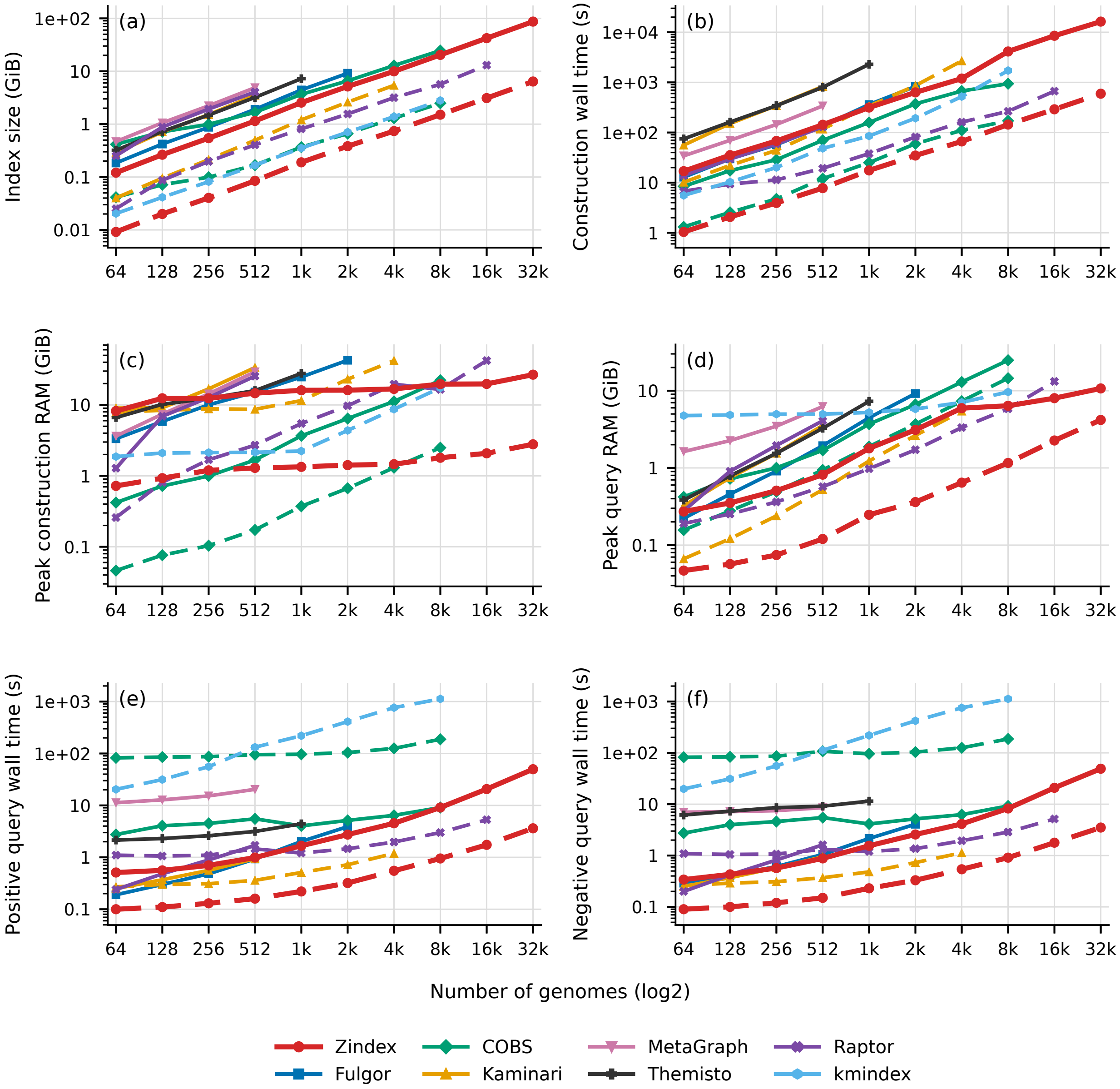


Fig. S1: RefSeq, target FPR $2^{-4}$. (a) Final index size; (b) construction CPU time; (c) construction wall time; (d) peak construction RSS; (e) positive and (f) negative query wall time. Solid lines denote dense indexing and dashed lines minimizers. One common legend applies to all panels. Only complete construction-and-query cases are shown; Fulgor, MetaGraph and Themisto retain dense curves only. Query times include loading and are medians of three repetitions. Shared external minimizer preparation is excluded; native OCM extraction is included.

### S2 RefSeq at 8-bit precision

Zindex uses native OCM31/window49. Other tools receive SIMD 31-mer minimizers, except historical ATB 8-bit Raptor, which uses its own 31/49 sampler. Raptor uses native window31 for supplied representatives. This reproduces the main RefSeq experiment with construction RAM and the two query classes shown separately. Zindex's sampled representation combines low storage and fast queries, while COBS requires less construction RAM at many shared sizes. The dense COBS query slowdown is I/O-bound and is retained.

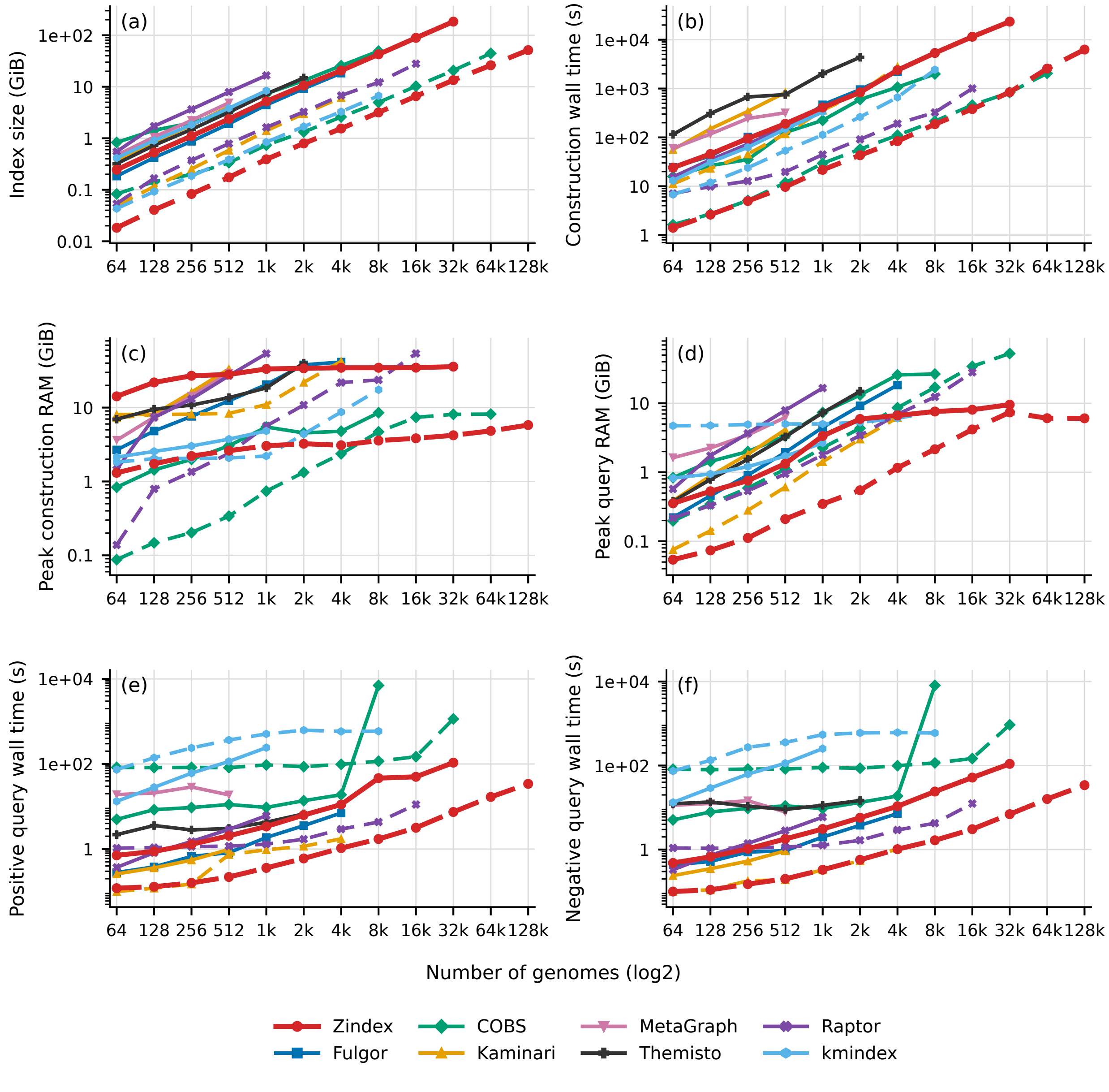


Fig. S2: RefSeq, target FPR $2^{-8}$. (a) Final index size; (b) construction CPU time; (c) construction wall time; (d) peak construction RSS; (e) positive and (f) negative query wall time. Solid lines denote dense indexing and dashed lines minimizers. One common legend applies to all panels. Only complete construction-and-query cases are shown; Fulgor, MetaGraph and Themisto retain dense curves only. Query times include loading and are medians of three repetitions. Shared external minimizer preparation is excluded; native OCM extraction is included.

### S3 RefSeq at 16-bit precision

Higher precision increases approximate-index storage while leaving exact indexes unchanged. Fulgor therefore becomes a stronger dense-space competitor, whereas Zindex remains compact with minimizers. Larger construction memory and intermediate-storage requirements limit Raptor and kmindex. Implementation differences across campaigns also contribute to comparisons with the historical 8-bit results.

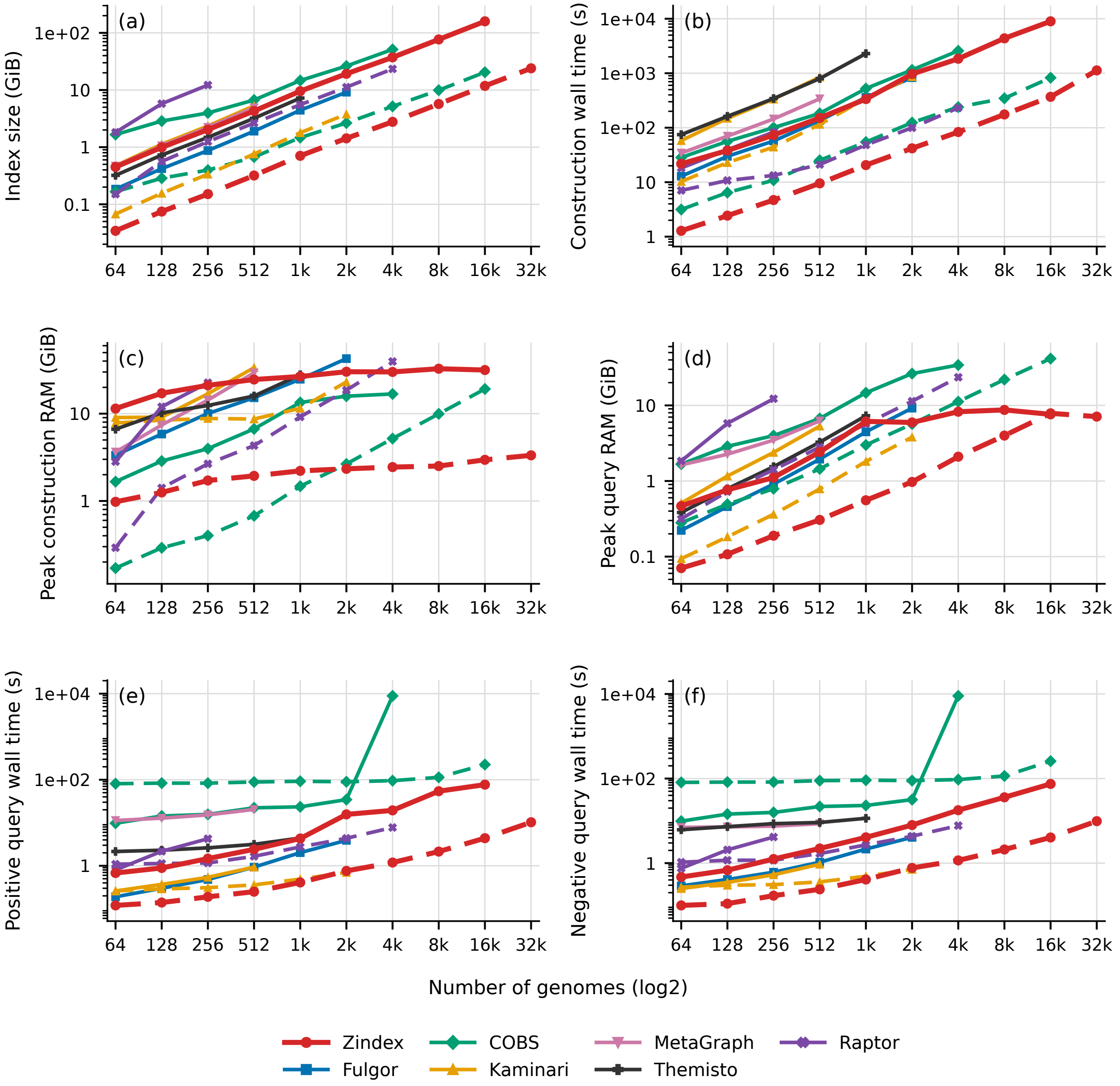


Fig. S3: RefSeq, target FPR $2^{-16}$. (a) Final index size; (b) construction CPU time; (c) construction wall time; (d) peak construction RSS; (e) positive and (f) negative query wall time. Solid lines denote dense indexing and dashed lines minimizers. One common legend applies to all panels. Only complete construction-and-query cases are shown; Fulgor, MetaGraph and Themisto retain dense curves only. Query times include loading and are medians of three repetitions. Shared external minimizer preparation is excluded; native OCM extraction is included.

### S4 ATB at 8-bit precision

Redundancy improves Zindex's footprint relative to diverse RefSeq, but benefits Fulgor's dense representation still more. Themisto and Kaminari also become more competitive in space. Zindex retains a compact sampled index and short construction times; its query advantage over COBS persists, while the comparison with other fast tools is less uniform.

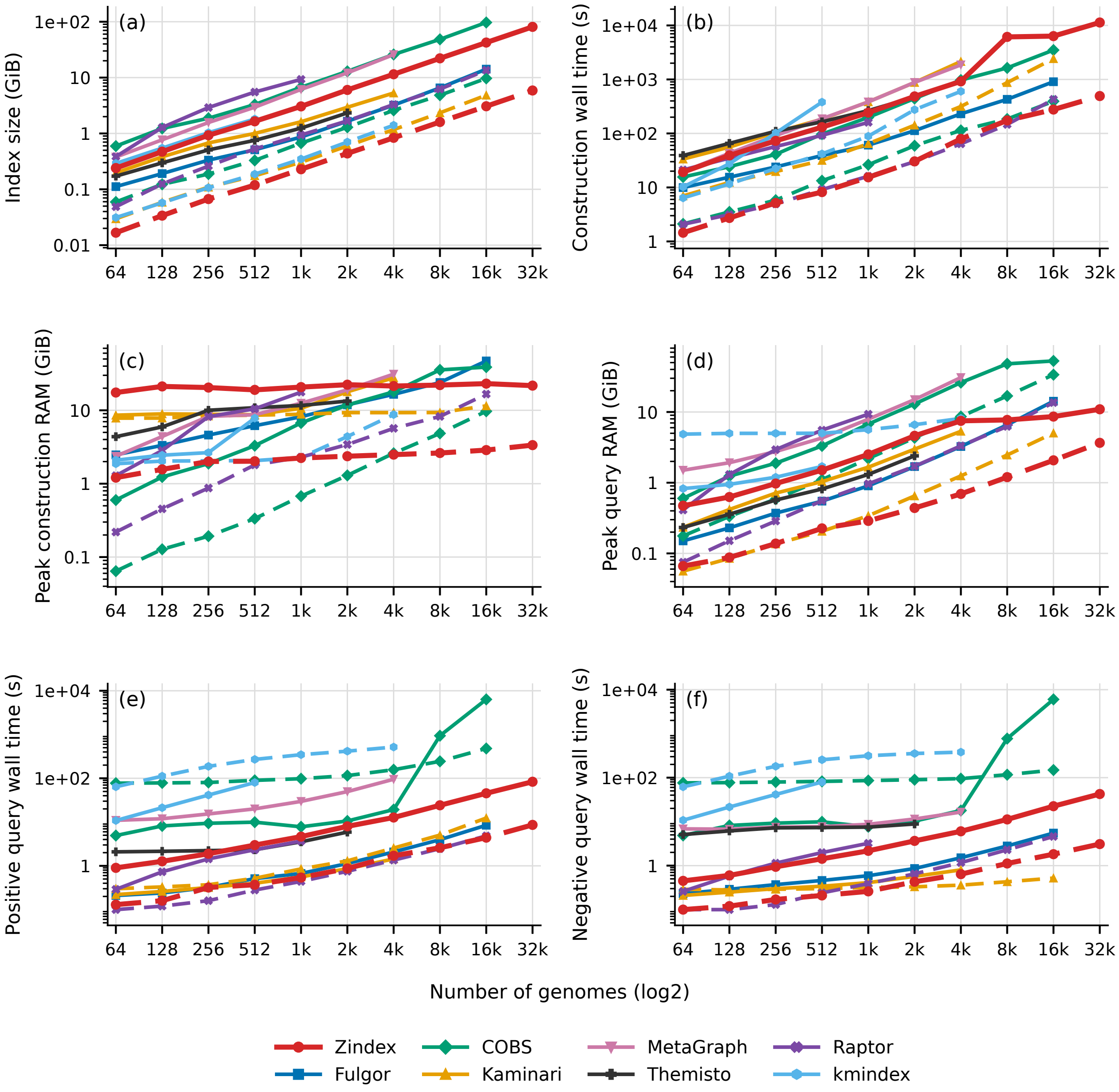


Fig. S4: ATB, target FPR $2^{-8}$. (a) Final index size; (b) construction CPU time; (c) construction wall time; (d) peak construction RSS; (e) positive and (f) negative query wall time. Solid lines denote dense indexing and dashed lines minimizers. One common legend applies to all panels. Only complete construction-and-query cases are shown; Fulgor, MetaGraph and Themisto retain dense curves only. Query times include loading and are medians of three repetitions. Shared external minimizer preparation is excluded; native OCM extraction is included.

### S5 ATB at 16-bit precision

At higher precision, Fulgor remains particularly compact in dense mode and Kaminari is a close sampled-space competitor to Zindex. Zindex's sampled construction and queries remain fast. This campaign is incomplete; its endpoints are not final scalability limits. ATB at 4-bit precision has no complete measurements in this snapshot.

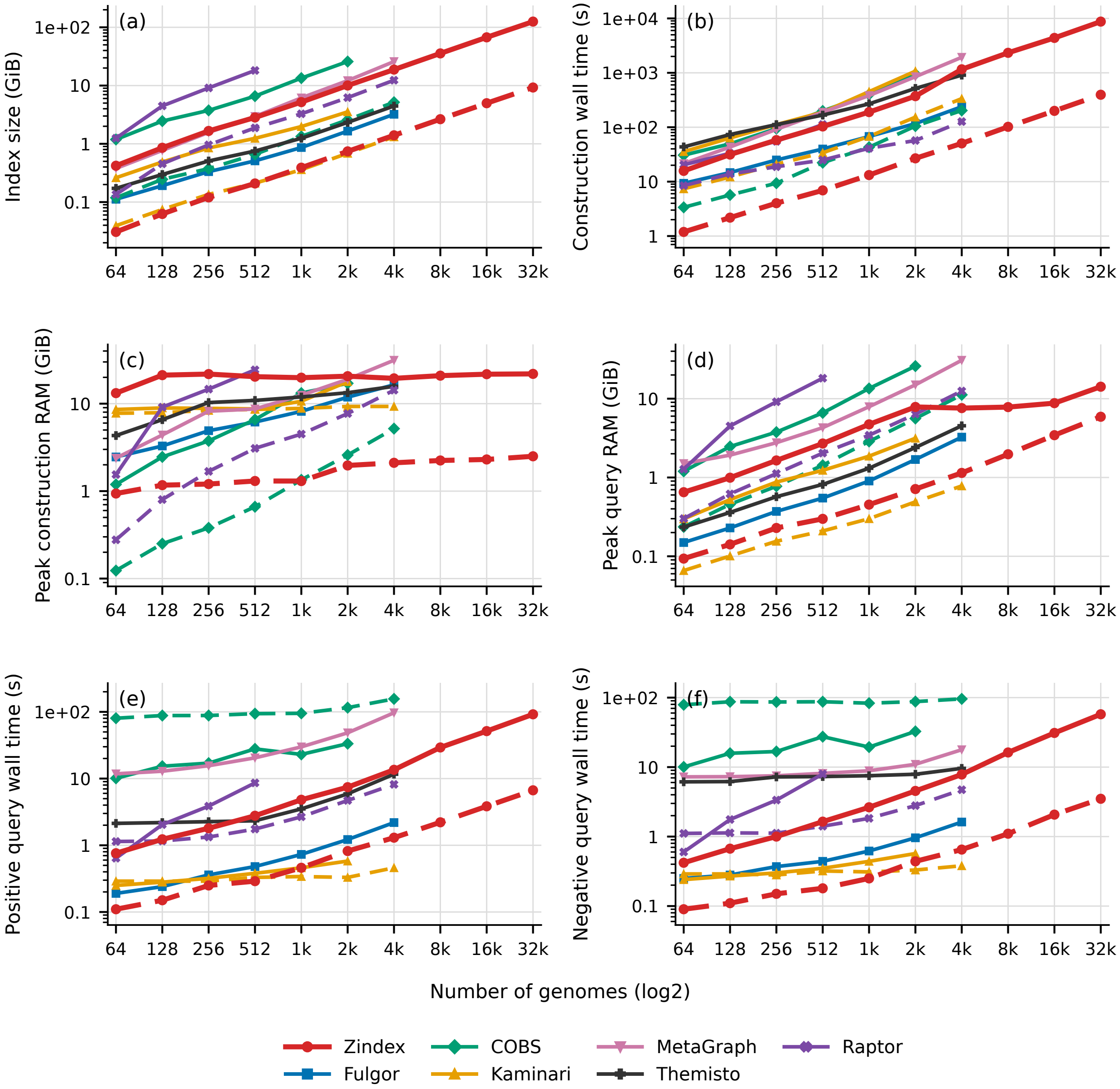


Fig. S5: ATB, target FPR $2^{-16}$. (a) Final index size; (b) construction CPU time; (c) construction wall time; (d) peak construction RSS; (e) positive and (f) negative query wall time. Solid lines denote dense indexing and dashed lines minimizers. One common legend applies to all panels. Only complete construction-and-query cases are shown; Fulgor, MetaGraph and Themisto retain dense curves only. Query times include loading and are medians of three repetitions. Shared external minimizer preparation is excluded; native OCM extraction is included.

### S6 Parameter sensitivity

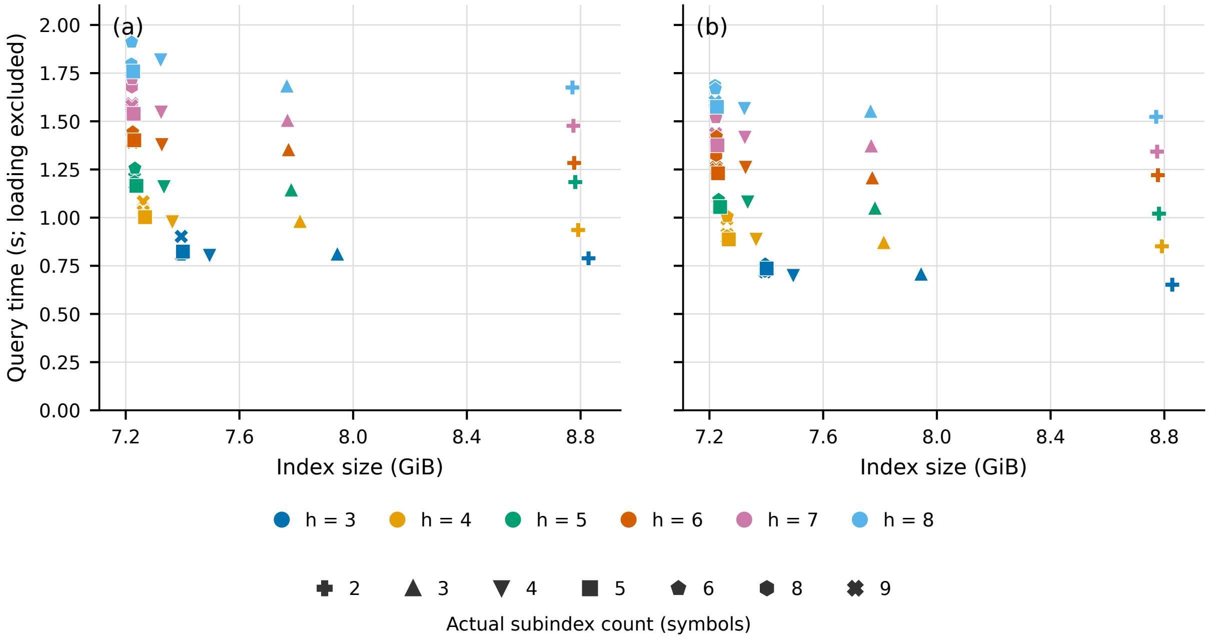


Fig. S6: Zindex parameter sensitivity on 16,384 bacterial genomes: (a) positive and (b) negative query time, excluding loading. Colors identify hash counts and symbols the actual subindex counts. The detail view omits one-subindex configurations and the seven-subindex series. Each point is an individual completed run.

### S7 Layout mechanisms

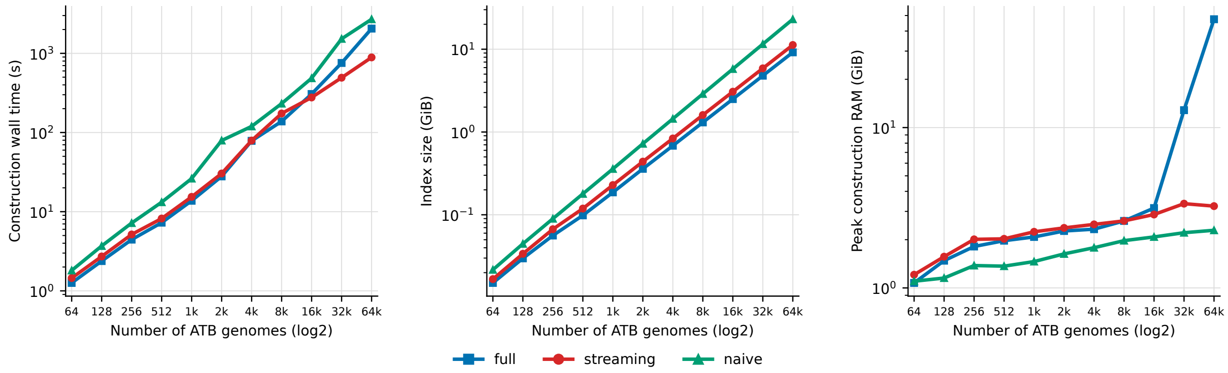


Fig. S7: Zindex construction strategies on ATB at target FPR $2^{-8}$. (a) Construction wall time; (b) final index size; (c) peak construction RSS. Full, streaming and naive use minimizers (k=31,w=19), eight hash functions and 32 threads. Full exceeded the 48 GB RAM limit at 65,536 genomes.

### S8 Large-scale runs and ablations

Table S1 lists every construction run on the two complete comparative collections, including the configurations that did not complete, and four single-parameter ablations of Zindex. Ordering the 661k input by lineage rather than by Raptor's input order reduced the index by 4.4% and construction time by 4%, supporting the taxonomic streaming order of Section 2.2. Increasing the number of hash locations from four to eight shrank the abandoned-key map by 19% while keeping interleaved filters the same size, but increased the index on disk by 16%: with more slots occupied, the filter arrays contain fewer zeros and compress less effectively. On *M. tuberculosis*, the 3% merge margin of Equation (2) reduced the overshoot of the actual bank size relative to the plan from 1.86 to 1.60 times, cut peak memory from 62.8 to 35.3 GB and shortened construction by 19%, at a cost of 3% in index size. Reducing the thread count from 32 to 8 produced an identical index, with construction taking 2.6 times longer while peak memory fell to 22.4 GB, so memory can be traded for time on smaller machines.

Table S1: Construction runs on the complete comparative collections, on M2 with minimizers of 19-mers from 31-base windows. Zindex defaults: open-closed mod-minimizers, merge margin 0.03, four hash locations, lineage order, 32 threads; each ablation changes one setting. Sizes in decimal GB; RAM is peak RSS. COBS wall time is minimizer extraction plus construction.

| Collection | Tool | Variant | Wall (h:mm) | RAM (GB) | Index (GB) |
|---|---|---|---|---|---|
| 661k | Raptor | default | out of memory at 2:46 | 803.6 | – |
| | Raptor | no rearrangement | >24:00 | – | – |
| | Raptor | no union estimation | >24:00 | – | – |
| | Zindex | Raptor input order | 11:49 | 33.8 | 101.5 |
| | Zindex | default (lineage order) | 11:22 | 33.1 | 97.0 |
| | Zindex | eight hash locations | 12:34 | 36.2 | 112.5 |
| *M. tuberculosis* | Raptor | default | >24:00 | – | – |
| | COBS | FPR 0.3, one hash | 24:40 | 30.2 | 100.3 |
| | Zindex | merge margin 0 | 1:20 | 62.8 | 48.8 |
| | Zindex | default | 1:05 | 35.3 | 50.5 |
| | Zindex | 8 threads | 2:46 | 22.4 | 50.5 |
| | Zindex | random minimizers | 2:36 | 65.0 | 71.7 |

Table S2: Direct construction versus index fusion for 32,768 RefSeq bacterial genomes, using native OCM minimizers ($k = 31$, $w = 19$), eight hash functions and 8-bit fingerprints. Input indexes were constructed sequentially, with all runs configured for 32 threads. Construction wall time includes input-index construction and fusion; fusion wall time is also reported separately. RAM denotes peak construction RSS. Query wall time combines the positive and negative workloads, each comprising 16,384 sequences of 1 kb, including loading, decompression, parsing and output. Sizes use decimal GB. † Includes approximately 91 minutes of host suspension.

| Input indexes | Index size (GB) | Construction (s) | Fusion (s) | Peak RAM (GB) | Query (s) |
|---|---|---|---|---|---|
| Direct | 14.42 | 881.90 | – | 4.62 | 14.91 |
| 2 × 16 384 | 14.37 | 921.15 | 38.99 | 4.51 | 15.88 |
| 4 × 8 192 | 14.29 | 947.34 | 42.59 | 4.33 | 17.32 |
| 8 × 4 096 | 14.25 | 1009.67 | 26.42 | 4.01 | 18.99 |
| 16 × 2 048 | 14.24 | 1084.05 | 30.24 | 3.81 | 20.23 |
| 32 × 1 024 | 14.20 | 959.41 | 20.60 | 3.97 | 19.31 |
| 64 × 512 | 14.16 | 994.92 | 40.27 | 3.74 | 21.85 |
| 128 × 256 | 14.10 | 902.23 | 33.25 | 3.64 | 22.21 |
| 256 × 128 | 14.06 | 966.15 | 18.68 | 3.55 | 26.19 |
| 512 × 64 | 14.05 | 1082.68 | 32.36 | 3.09 | 33.13 |

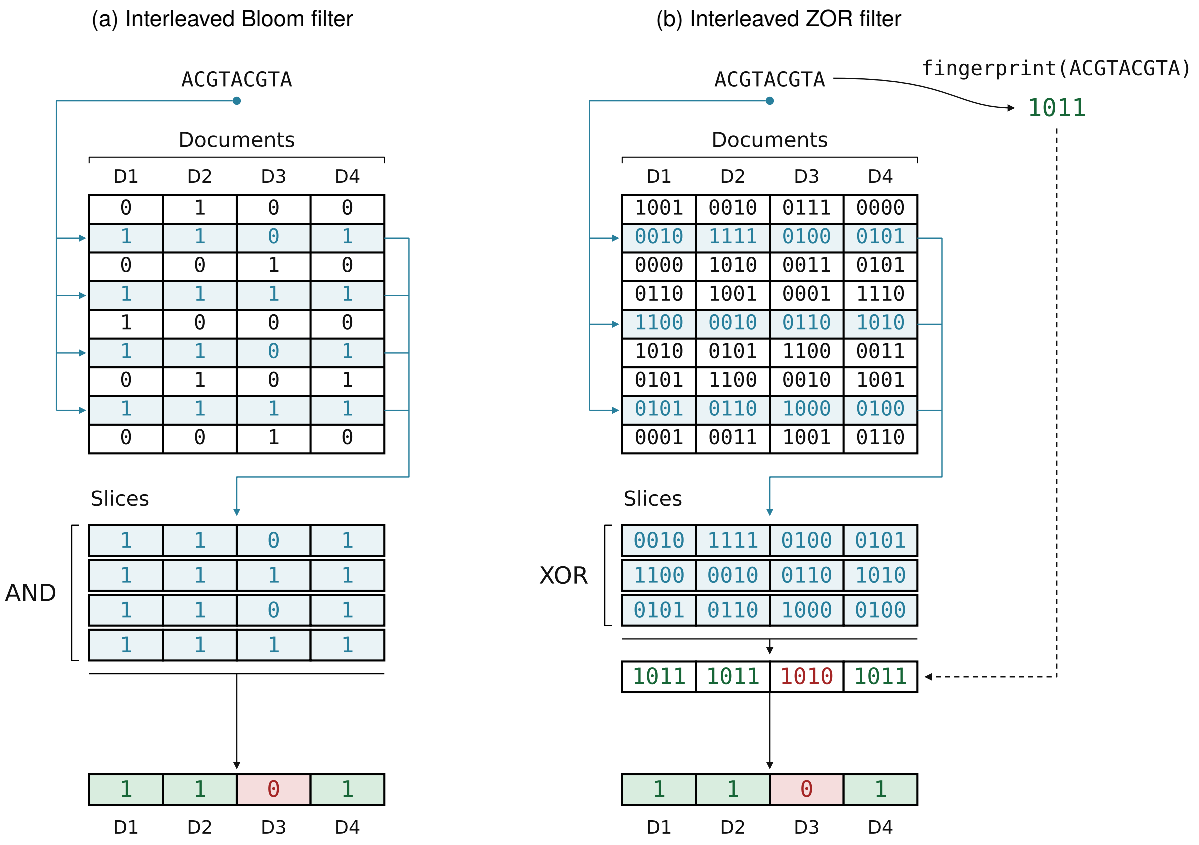


Fig. S8: Interleaved filters over four documents, queried with one $k$-mer. (a) Interleaved Bloom filter. The four hash functions select four rows, and the slices are ANDed column-wise; a document is reported only if all four bits are 1, so $D_3$ is rejected. (b) Interleaved ZOR filter with 4-bit fingerprints. The three hash functions select three rows, the slices are XORed column-wise, and each document's result is compared with the query fingerprint $F(\texttt{ACGTACGTA}) = \texttt{1011}$; $D_3$ yields `1010` and is rejected. In both layouts, one pass over a few contiguous rows answers the query for every document at once, but the Bloom filter needs more rows and more bits per key for the same false-positive rate.
